# Heather pollen is not necessarily a healthy diet for bumble bees

**DOI:** 10.1101/2023.04.06.535809

**Authors:** Clément Tourbez, Irène Semay, Apolline Michel, Denis Michez, Pascal Gerbaux, Antoine Gekière, Maryse Vanderplanck

## Abstract

There is evidence that specialised metabolites of flowering plants occur in both vegetative parts and floral resources (i.e., pollen and nectar), exposing pollinators to their biological activities. While such metabolites may be toxic to bees, it may also help them to deal with environmental stressors. One example is heather nectar which has been shown to limit bumble bee infection by a trypanosomatid parasite, *Crithidia* sp., because of callunene activity. Besides in nectar, heather harbours high content of specialised metabolites in pollen such as flavonoids but they have been poorly investigated. In this study, we aimed to assess the impact of *Crithidia* sp., heather pollen and its flavonoids on bumble bees using non-parasitised and parasitised microcolonies fed either control pollen diet (i.e., willow pollen), heather pollen diet, or flavonoid-supplemented pollen diet. We found that heather pollen and its flavonoids significantly affected microcolonies by decreasing pollen collection as well as offspring production, and by increasing male fat body content while parasite exposure had no significant effect except for an increase in male fat body. We did not highlight any medicinal effect of heather pollen or its flavonoids on parasitised bumble bees. Our results provide insight into the impact of pollen specialised metabolites in heather-bumble bee-parasite interactions. They underline the contrasting roles for bumble bees of the two floral resources and highlight the importance of considering both nectar and pollen when addressing medicinal effects of a plant towards pollinators.

## Introduction

For their own subsistence and that of their offsprings, bee females mostly forage on two floral resources, namely nectar as main source of carbohydrates (Nicolson & Thornburg, 2007), and pollen as main source of proteins and lipids (Campos et al., 2008). Among these nutritional resources, the pollen chemical composition is particularly complex and highly variable among plant species (Vaudo et al., 2020). While pollen central metabolites, for instance the protein-to-lipid ratio, play crucial role in bee health, development, and fitness (Di Pasquale et al., 2013), pollen also contains numerous specialised metabolites (e.g., alkaloids, flavonoids and terpenoids, Irwin et al., 2014; Palmer-Young et al., 2019). The biological activities of these metabolites are multiple so that they may be involved in protecting pollen from abiotic factors, such as UVs (Li et al., 1993), but also from biotic factors, acting as antibacterial, antifungal or insecticidal compounds (Pusztahelyi et al., 2015; Zaynab et al., 2018). When ingesting pollen, bees are then exposed to all these biological activities that may be beneficial, for instance by reducing parasite load through antimicrobial activities (Manson et al., 2010; Biller et al., 2015; Richardson et al., 2015), but also detrimental, for instance by impairing resource collection (Wang et al., 2019; Brochu et al., 2020), decreasing offspring size and production (Arnold et al., 2014), inducing larvae or imago death (Hendriksma et al., 2011; Weber, 2004), and altering immune system (Gekière et al., 2022a). Given these opposite effects on bees, it is essential to question how specific specialised metabolites may impact bee health, especially in a changing world with multiple environmental pressures.

In the current context of biodiversity erosion (Butchart et al., 2010), bees are unfortunately not exception, and many threats have been pinpointed as responsible for their negative population trends (Dicks et al., 2021) such as pesticide exposure (Sánchez-Bayo & Goka, 2014), metalloid pollution (Gekière et al., 2023), habitat loss (Baude et al., 2016), resource scarcity (Naug, 2009), competition with domesticated species (Mallinger et al., 2017), and diseases (Van Engelsdorp et al., 2009). Among environmental challenges, bees indeed suffer from a high diversity of pathogens and parasites (Meeus et al., 2011; Goulson & Hughes, 2015) of which effects vary from small ethological alterations of minor consequences (Paris et al., 2018) to large reductions in host bee fitness (McMenamin & Genersch, 2015). Social bee species such as bumble bees (Apidae; *Bombus* spp.) are particularly impacted by parasites, the latter benefiting from their social system to readily infect numerous individuals (Folly et al., 2017). One of the most prevalent parasites in wild bumble bee populations is the gut trypanosomatid *Crithidia bombi* Lipa & Triggiani, 1980 (Euglenozoa: Trypanosomatidae; Schmid-Hempel, 2001). Despite its generally moderate impacts, it can decrease foraging effectiveness (Otterstatter et al., 2005), offspring production (Schmid-Hempel, 1998), queen survival through hibernation (Fauser et al., 2017), and increase mortality in synergy with other stresses (Brown et al., 2000). To deal with such parasite pressure, bumble bees may rely on specific floral resources displaying appropriate antimicrobial properties through their specialised metabolites (Manson et al., 2010; Biller et al., 2015; Richardson et al., 2015; Fitch et al., 2022).

Among potential medicinal floral resources, heather (Ericaceae, *Calluna vulgaris* Hull. 1808) produces a nectar documented to affect *C. bombi* (Koch et al., 2019). This effect has been attributed to the presence of callunene, a terpenoid that induces the loss of *C. bombi* flagellum, preventing the parasite from settling in the bumble bee digestive tract (Koch et al., 2019). Such medicative properties of heather nectar make heather-rich heathlands even more valuable for these bumble bees (Descamps et al., 2015; Moquet et al., 2017). However, although heather is a major resource for European bees, only a handful of studies have sought for specialised metabolites with biological activities in heather pollen, showing a high prevalence of flavonoids (Gekière et al., *in prep*.). Flavonoids can have very contrasting effects on insect-plant interactions and affect them in multiple ways (Simmonds 2003; Onyilagha et al., 2012). Bees are attracted to some flavonoid compounds (e.g. quercetin; Liao et al. 2017a) while others repel them (e.g. kaempferol, catechin; Detzel & Wink 1993; Onkokesung et al., 2014). However, despite some deleterious effects on larval development (Wang et al., 2010), flavonoids are mainly not toxic molecules for insects (Detzel & Wink, 1993). Once ingested, flavonoids can have antioxidant properties and are potentially beneficial for bees (e.g. quercetin; Treutter, 2005). They can stimulate the activation of detoxification enzymes (cytochrome P450 monooxygenase) and enhance bee resistance to certain insecticides and acaricides (Scott et al., 1998; Johnson et al. 2012; Liao et al., 2017b). The case of heather pollen flavonoids remains to be adressed and this incomplete picture of the pollen side does not allow for fully arguing that heather is a bumble bee health-promoting plant. Therefore, bioassays to determine heather pollen effects on bumble bee brood, bumble bee health, and parasite dynamics are warranted. To fill this gap, we herein present a study that aimed to assess the effects on heather pollen and its flavonoids on bumble bee health, at both individual and colony levels, considering the bumble bee interaction with the parasite. We specifically addressed the following questions: (i) how does the parasite influence the development of bumble bee microcolonies and the individual immunocompetence? (ii) do heather pollen and its flavonoids challenge bumble bees, impacting their resource collection and offspring production? (iii) do heather pollen and its flavonoids affect the parasite dynamic in infected bumble bee workers, or help bumble bees to counteract parasite effects? We expect (i) a mild effect of the parasite on bumble bees reared in optimal conditions; (ii) detrimental effects of flavonoids, and potentially of heather pollen on healthy bumble bees and microcolonies; and (iii) beneficial effects of heather pollen, and potentially its flavonoids, on infected bumble bees by reducing parasite load.

## Materials and methods

### Bumble bee bioassays

Queenless microcolonies of five workers were exposed to specific diet treatments (Fig.1): control pollen (i.e., willow pollen is used because artificial pollen is unsuitable for bumblebee development and because its flavonoid profile does not overlap with any flavonoids found in heather pollen; Gekière et al., 2022b; Gekière et al., *in prep*) containing either (i) parasitised or (ii) non-parasitised bumble bees; heather pollen containing either (iii) parasitised or (iv) non-parasitised bumble bees; microcolonies fed with willow pollen supplemented with extracts of flavonoids from heather pollen containing either (v) parasitised or (vi) non-parasitised bumble bees. Diets (i) and (ii) were used as controls as well as to assess the parasite impacts. Diets (iii) and (v) were used to establish the effects of heather pollen or its flavonoids on infected microcolonies. Diets (iv) and (vi) were used to establish the effects of heather pollen or its flavonoids on uninfected microcolonies. For each treatment, ten microcolonies have been established using five different queenright colonies (colonies from Biobest *bvba*; Westerlo, Belgium) (2 microcolonies per colony per treatment). Faeces of queenright colonies were observed under the microscope to confirm the absence of parasites (*Nosema* spp., *Apicystis* spp. and *Crithidia* spp.) as guaranteed by the supplier. The microcolonies were kept in plastic boxes (10 × 16 × 16 cm; Regali & Rasmont, 1995) and reared at the University of Mons (Belgium, Mons, Campus de Nimy, WGS84 50°27’54.9’’N 3°57’24.9’’E) in a dark room at 26-28°C and 65% of relative humidity. Bumble bees were provided *ad libitum* with sugar syrup (water/sugar 35:65 w/w) and pollen candies (i.e., pollen mixed with a 65% sugar solution) for 35 days, with pollen candies being freshly prepared and renewed every two days. When workers died, they were discarded, weighted and replaced by a worker from the same queenright colony, which was marked with a colour dot on the scutum. Larvae ejected from the brood were also checked every day, counted and discarded from the microcolonies. Microcolonies were handled under red right to minimise disturbance.

**Figure 1.**
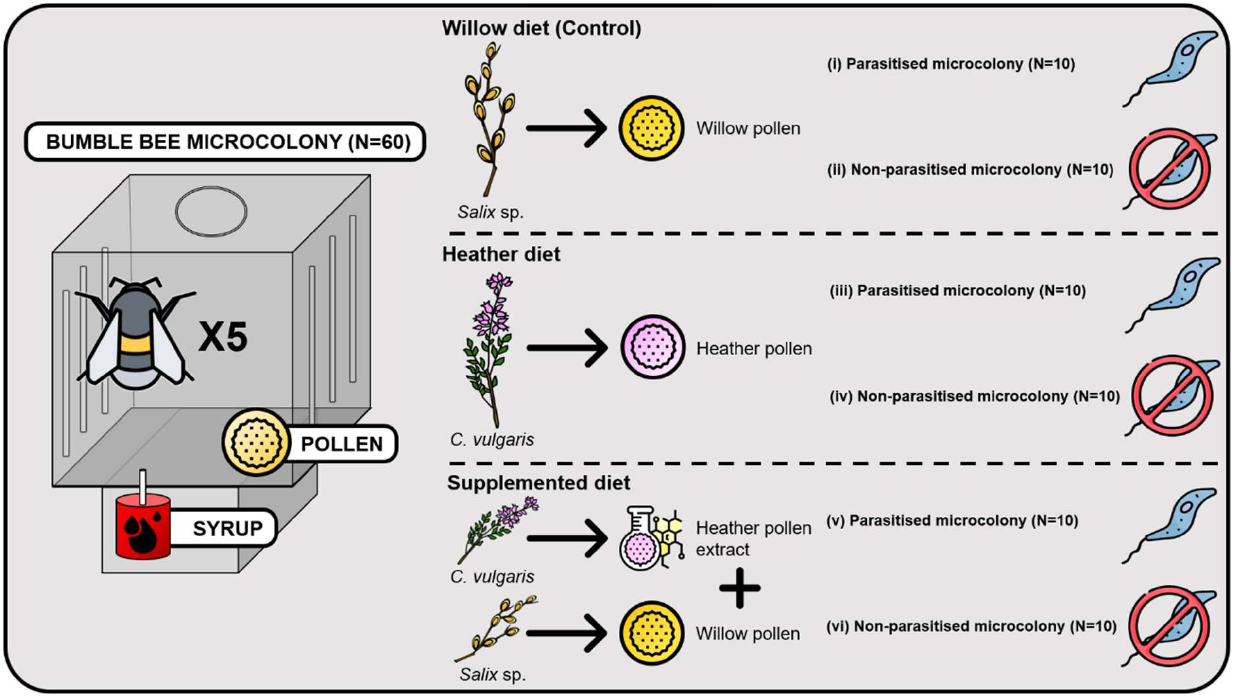
Bioassay design. Microcolonies initiated with five *B. terrestris* workers were fed for 35 days with one of three diets. For each diet, ten microcolonies contained parasitised individuals (*Crithidia* sp.), ten others were non-parasitised. Icon used for the figure: https://www.flaticon.com/. and author conception.

### Diet preparation

Willow pollen batch (i.e., pollen loads from *Apis mellifera* L. 1758) was supplied by the commercial company *Ruchers de Lorraine* (Nancy, France) while heather pollen batch was obtained from a private beekeeper (Dittlo François, France, Gironde, Le Nizan). Although honey bee collected pollen loads may contain parasite, analysis of faeces of uninfected microcolonies fed this pollen diet was parasite free. We then consider that no contamination occurred from the pollen batch. Pollen loads from the heather batch were hand-sorted based on the colour after microscopical identification to ensure monoflorality (800 g in total) (Sawyer & Pickard, 1981; Dafni et al., 2005). Each pollen batch was then homogenised and crushed before being used for the experiments. Half of the sorted heather batch served directly for the bioassays; the other half served for massive extraction of flavonoids. Flavonoids were extracted using a Soxhlet extraction for approximatively 40 cycles with methanol as solvent, at 100°C. Extract was then vacuum filtered and evaporated to dryness (rotavapor IKA RV8). For flavonoid purification, extract was solubilized in water with a minimal amount of methanol, and placed in a separatory funnel with dichloromethane. The funnel was shaken and left to settle overnight before recovering the aqueous phase. The purified extract was then dried using rotary evaporator and dissolved in aqueous ethanol solution (1:1, v/v) before addition to the control diet to prepare a flavonoid supplemented diet. Control and heather pollen diets were also supplemented with a similar amount of ethanol to avoid any bias (for details see Appendix A, Table S1).

### Parasite inoculation

Multiple morphologically identical trypanosomes affect *B. terrestris* (Bartolomé et al., 2021). Although *Crithidia bombi* is by far the most abundant in wild populations (Shykoff & Schmid-Hempel, 1991; Popp et al., 2012), parasite identification will be limited to *Crithidia* sp. in this manuscript to avoid misinterpretation. Parasite inoculation was performed using *Crithidia* sp. reservoirs maintained in the laboratory (i.e., commercial colonies regularly renewed and repeatedly inoculated with contaminated faeces in order to ensure a turnover of the available *Crithidia* sp. pool). Faeces from a total of 45 infected workers were collected and pooled together to ensure multiple-strain inoculum (Gekière et al., 2022a). The inoculum was homogenised, brought to 1 mL with 0.9% NaCl solution, and purified by a triangulation method (Cole, 1970) adapted by Baron et al. (2014) and Martin et al. (2018). The concentration of *Crithidia* sp. cells was then estimated by counting with a Neubauer chamber, and the inoculum was diluted to 2,500 *Crithidia* sp. cells/µL with a 40% sugar solution. Workers allocated to the infected microcolonies were placed in individual Nicot® queen rearing cages and given 10µL of the inoculum (i.e., 25,000 *Crithidia* sp. cells; Logan et al., 2005) by letting them feed on the sugar solution in a glass microcapillary after a 5-hour starvation period. Workers allocated to uninfected microcolonies also underwent the same treatment (isolation, starvation) but with 10µL of sterile sugar solution.

### Parameters evaluated

To investigate the impacts of pollen diet and parasite, several parameters in microcolonies were measured (Tasei & Aupinel, 2008), namely resource collection, reproductive success, stress response, and individual health through fat body content (i.e., immunocompetence proxy; Arrese & Soulages, 2010; Rosales, 2017; Vanderplanck et al., 2021) and parasite load measurements.

Resource collection was assessed by weighting every two days in each microcolony the syrup container, as well as the recovered pollen candy and the newly introduced one. These data were corrected for evaporation using controls, as well as divided by the total worker mass by microcolony to avoid bias due to worker activity. To evaluate the reproductive success, all microcolonies were dissected at the end of the experiment (Day 35) to weigh the total hatched brood mass, as well as the individual mass of each emerged male used as reference of viable offspring at the end of development (Goulson, 2010). Offspring masses were divided by the total worker mass by microcolony to avoid any bias due to worker care. Regarding stress response, we assessed worker mortality, larval ejection, pollen dilution (ratio between the collection of syrup and pollen) as well as pollen efficiency (ratio between offspring mass and pollen collection; Tasei & Aupinel, 2008) that highlights when a micro-colony needs to consume more pollen to produce offspring.

For the individual health parameters, fat body content was measured at the end of the bioassays on two males and two workers per microcolony (40 individuals per treatment) following Ellers (1996). The abdomens were cut and dehydrated in an incubator at 70°C for three days before being weighed. They were then placed for one day in 2mL of diethyl ether to solubilise lipids constituting the fat body. The abdomens were then washed twice with diethyl ether, and incubated at 70°C for seven days before being weighed. Fat body content was defined as the mass difference between dry abdomen before and after lipid solubilisation, divided by the dry abdomen mass prior to solubilisation. Moreover, in infected treatments, we repeatedly monitored the parasite load within microcolonies using the same marked worker along the bioassays. The first measurement was made three days post-inoculation (day 4) to enable *Crithidia* sp. to multiply, and ensure its presence in the faeces (Logan et al., 2005). A total of seven further measurements were taken to establish the infection curve of *Crithidia* sp., namely at days 6, 8, 10, 12, 16, 20 and 35. Measurements were performed at larger intervals after day 12 because infection reached the plateau phase (Schmid-Hempel & Schmid-Hempel, 1993; Otterstatter & Thomson, 2006). In practice, the marked worker was held in a 50mL Falcon in the light until the faeces were expelled. Faeces were then collected in a 10µL microcapillary tube and diluted two to ten times with distilled water to enable rational cell counting. Parasite cells were then counted using a haemocytometer (Neubauer) under an inverted phase contrast microscope (400X magnification, Eclipse Ts2R, Nikon). Uninfected microcolonies faeces were checked to be free of parasite at the end of the experiment, and a marked worker was isolated at days 4, 6, 8, 10, 12, 16, 20 and 35 in each uninfected microcolony to induce the same stress as in infected treatments.

### Data analysis and statistics

To detect a potential effect of pollen diet or parasite on resource collection, reproductive success, stress response, and individual health, mixed models were fitted for each parameter using diet, parasite and their interaction as fixed factors, and colony as random factor. Pollen collection, pollen efficacy, and pollen dilution (log-transformed data) were analysed using a Gaussian distribution (i.e., normality of residuals; *shapiro*.*test* function from the stats R-package v.4.1.0; R Core Team, 2021) (*lmer* function from the nlme R package v.3.1.160; Pinheiro et al., 2022). Total hatched offspring mass, emerged male mass, and fat body content (i.e., proportion data) were analysed using a Gamma distribution and a log link function (*glmmTMB* function from the glmmTMB R-package v.1.1.4; Brooks et al., 2017). For fat body content, values were square root-transformed and sex was added as crossed fixed effect. For emerged male mass and fat body content, microcolony was nested within colony in the random factor to deal with pseudo-replication (i.e., several measures per microcolony).

For larval ejection, a binomial distribution (ejected larvae and total number of living offspring produced as bivariate response) was used after checking for overdispersion and zero inflation (*testDispersion* and *testZeroInflation* functions from DHARMa R-package v.0.4.6; Hartig, 2022). For worker mortality, a Cox proportional hazard (mixed-effect) model was run with individuals alive at the end of the 35-day treatment assigned as censored, and those who died as uncensored (*coxme* function from the coxme R-package v.2.2.18.1; Therneau, 2022).

The last parameter measured was the parasite load at different time points within infected microcolonies. As infection dynamics is a discrete time series, it was analysed using a generalised additive mixed-effect model (GAMM; Wood, 2006). Parasite load were square root-transformed and fitted using a Gaussian distribution with a log link. Diet and day were set as fixed factor and microcolony nested within colony as random factor. The model assumptions were tested using diagnostic graphs and tests.

Contrasts were then performed on the models to determine whether uninfected control differed from the infected control, whether effects on uninfected or infected microcolonies differed among diets (*emmeans* function from the emmeans R-package v.1.8.2; Lenth, 2022). For fat body content, data were analysed separately for workers and males as a sex significant effect was detected. Graphs and plots were all performed using the R-package ggplot2 v.3.4.0 (Wickham, 2016) except the one referring to the survival probability of the workers performed with the *ggsurvplot* function of the survminer R-package v.0.4.9 (Kassambara et al., 2021). All the statistical analyses were done using the R software v.4.1.0 (R Core Team, 2021). For all statistical analyses, *p* < 0.05 was used as a threshold for significance.

## Results

### Parasite impact

Comparison of microcolonies fed with control pollen between parasitised and non-parasitised treatments showed that *Crithidia* sp. infection did not impact the parameters related to resource collection (Fig 2A), reproductive success (Fig 2B-2C), or stress response (Fig 3A-C) (*p >* 0.05, Fig. 2 and 3). However, fat body content in newly emerged males was significantly higher with a mean that increased by 56% in infected microcolonies fed the control diet compared to uninfected ones fed the same diet (*t* = -3.828, *p* = 0.0012; Fig. 4B). The estimates (mean ± standard error) of our variables for each treatment are available in the appendices (Appendix B, Table S2)

**Figure 2.**
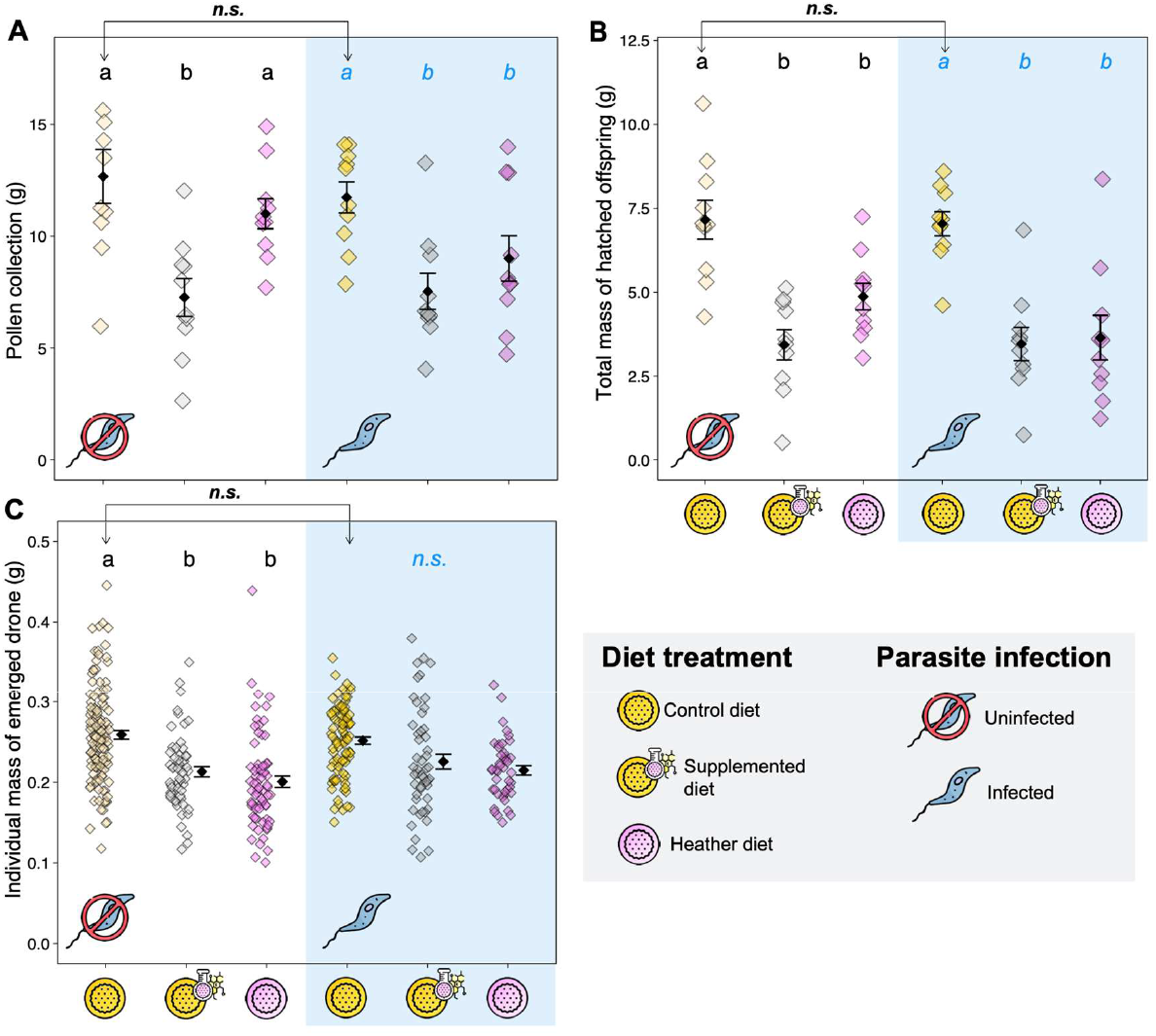
Resource collection and reproductive success. **(A)** Total mass of collected pollen; **(B)** Total mass of hatched offspring produced; and **(C)** Individual mass of emerged males. Each coloured data point represents a microcolony (in A and B) or an individual (in C), diamonds are mean values of each treatment, and error bars indicate the standard error. For **(C)**, means and error bars have been shifted in the graph to improve readability. Letters indicate significance at *p <* 0.05 (pairwise comparisons within uninfected treatments in black, and pairwise comparisons within infected treatments in blue); n.s., non-significant. Arrows indicate the pairwise comparisons for the control diet between infection treatments (i.e., parasite effect). Symbol caption is in the grey zone.

**Figure 3.**
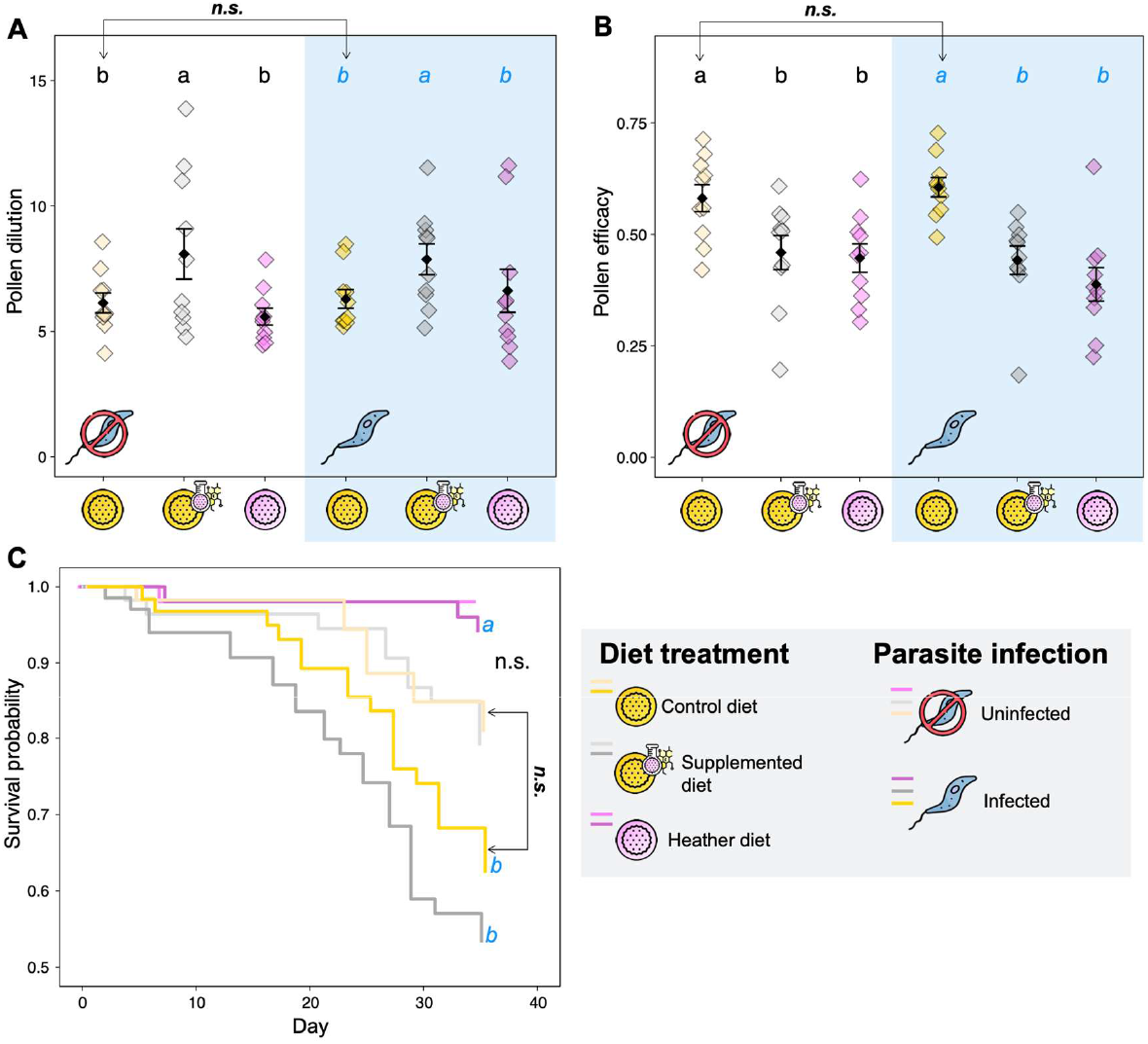
Stress responses. **(A)** Pollen dilution, defined as the ratio between syrup and pollen collection; **(B)** Pollen efficacy, defined as the ratio between total mass of hatched offspring and pollen collection; and **(C)** Worker survival probability over time. For **(A)** and **(B)**, each coloured data point represents a microcolony, diamonds are mean values of each treatment, and error bars indicate the standard error. Letters indicate significance at *p <* 0.05 (pairwise comparisons within uninfected treatments in black, and pairwise comparisons within infected treatments in blue); n.s., non-significant. Arrows indicate the pairwise comparisons for the control diet between infection treatments (i.e., parasite effect). Symbol caption is in the grey zone.

**Figure 4.**
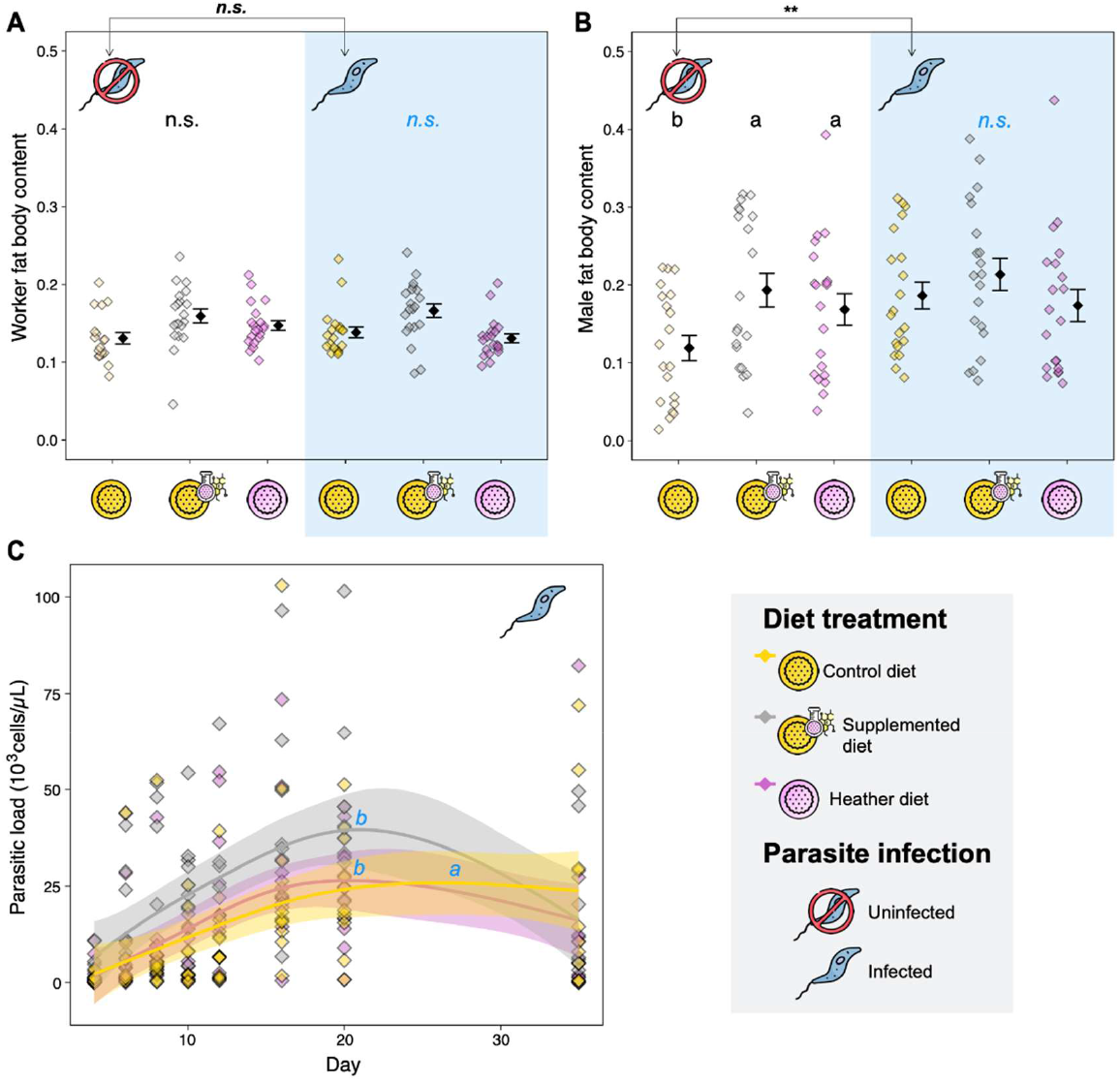
Health parameters. **(A)** Worker fat body content, and **(B)** Male fat body content; each coloured data point represents a microcolony, diamonds are mean values of each treatment, and error bars indicate the standard error. Means and error bars have been shifted in the graphs to improve readability. **(C)** Parasite load over time. Generalized additive mixed-effect models (in C) were used to fit smoothers to the data showing mean trends [±95 % confidence intervals, light coloured bands] over time. Here, each dot represents one data point (i.e., parasite load for the monitored worker for each time point and each microcolony). Letters indicate significance at *p <* 0.05 (pairwise comparisons within uninfected treatments in black, and pairwise comparisons within infected treatments in blue); n.s., non-significant; **, *p <* 0.01. Arrows indicate the pairwise comparisons for the control diet between infection treatments (i.e., parasite effect). Symbol caption is in the grey zone.

### Effect of heather pollen and its flavonoids on healthy bumble bees

Regarding resource collection, total pollen collection was significantly lower in microcolonies fed the supplemented diet compared to those fed the other diets (control *vs* supplemented: 43% less pollen collected, *t* = -5.672, *p* <0.001; heather *vs* supplemented: 33% less pollen collected, *t* = 3.924, *p* < 0.001; Fig. 2A). With regards to the reproductive success, microcolonies fed the supplemented and heather diets produced a significantly lower brood mass compared to microcolonies fed the control diet (control *vs* supplemented: brood mass 52% lower, *t* = 3.890, *p* < 0.001; control *vs* heather: brood mass 32% lower, *t* = 2.189, *p* = 0.0331; Fig. 2B), as well as significantly smaller emerged males (control *vs* supplemented: *t* = 2.350, *p* = 0.0192; control *vs* heather: *t* = 2.925, *p* = 0.0036; Fig. 2C)

In terms of stress responses, pollen dilution was significantly higher in microcolonies fed the supplemented diet compared to those fed the other diets (control *vs* supplemented: *t* = 2.282, *p* = 0.0268; heather *vs* supplemented: *t* = -3.191, *p* = 0.0025; Fig. 3A). Microcolonies fed the heather or supplemented diets also displayed a lower pollen efficacy than the microcolonies fed the control diet (control *vs* supplemented *t* = -2.741, *p* = 0.0085; control *vs* heather: *t* = -3.025, *p* = 0.0039; Fig. 3B). On the contrary, no significant difference was detected for larval ejection (*p >* 0.05) or for worker mortality (p < 0.05, Fig. 3C).

Regarding individual health, while no difference was detected in worker fat body content among diet treatments (*p >* 0.05; Fig. 4A), fat body content in newly emerged males was significantly higher in microcolonies fed the supplemented or heather diets compared to those fed the control diet (control *vs* supplemented: fat body content 62% higher, *t* = -3.891, *p* = 0.0012; control *vs* heather: fat body content 41% higher, *t* = 2.850, *p* = 0.0223; Fig. 4B).

### Effect of heather pollen and its flavonoids on parasitised bumble bees

Similarly as previous results with uninfected microcolonies, total pollen collection was significantly lower in infected microcolonies fed the supplemented diet than in microcolonies fed the control diet (36% less pollen collected, *t =* -4.414, *p <* 0.001), but also in infected microcolonies fed heather diet compared to those fed the control diet (16% less pollen collected, t = -2.866, p = 0.0061) (Fig. 2A). Regarding the reproductive success, as observed in uninfected microcolonies, microcolonies fed the supplemented and heather diets produced a significantly lower brood mass compared to microcolonies fed the control diet (control *vs* supplemented: brood mass 51% lower, *t* = 3.784, *p* < 0.001; control *vs* heather: brood mass 41% lower, *t* = 3.551, *p* < 0.001; Fig. 2B). However, no significant difference was detected for the mass of newly emerged males among diet treatments (*p <* 0.05; Fig. 2C).

Regarding stress responses, pollen dilution was significantly higher in microcolonies fed the supplemented diet than in microcolonies fed the other diets (control *vs* supplemented: *t* = 2.111, *p* = 0.0398; heather *vs* supplemented: *t* = -2.120, *p* = 0.0390; Fig. 3A). Microcolonies fed the heather or supplemented diets also displayed a lower pollen efficacy than those fed the control diet (control *vs* supplemented: *t* = -3.684, *p* < 0.001; control *vs* heather: *t* = -4.904, *p* < 0.001; Fig. 3B). While no significant difference was detected for larval ejection (*p >* 0.05), the worker survival probability was significantly reduced in infected microcolonies fed the heather diet compared to those fed either the control or supplemented diets (heather *vs* control: *t* = -2.265, *p* = 0.0235; heather *vs* supplemented: *t* = -3.331, *p* < 0.001; Fig. 3C).

In terms of individual health, no difference was detected in fat body content of workers or newly emerged males among diet treatments (*p >* 0.05; Figs. 4A and 4B). Regarding the parasite load, infection dynamic was more gradual in infected microcolonies fed the control diet compared to those fed the other diets which supported a parasite load peak around day 20 before a decrease up to the end of treatment (supplemented *vs* control: *t =* 2.893, *p =* 0.0126; heather *vs* control: *t =* 2.328, *p =* 0.0313; Fig. 4C).

## Discussion

### Parasite effect on bumble bee

The parasite had no impact neither on larval ejection, total mass of offspring produced, nor on individual mass of newly emerged males. Such results suggest that infection is unlikely to reduce colony and offspring fitness, or dissemination success, which are related to individual size (Greenleaf et al., 2007; Amin et al., 2012). The limited effects of *Crithidia* sp. on the reproductive success of bumble bees herein observed are in line with the literature (Brown et al., 2003; Goulson et al., 2018; Gekière et al., 2022a). This absence of impacts on development performance and offspring fitness may stem from the fact that the parasite only infects the adult stage (i.e., *Crithidia* sp. does not develop in bumble bee larvae, Folly et al., 2017).

Besides our results showed that *Crithidia* sp. induced larger fat body content in males emerged in infected microcolonies compared to those emerged in uninfected ones, whereas it has not impact on worker fat body content. We propose two rationales to explain such a *Crithidia*-induced difference in fat body content only in newly emerged males and not in workers. First, newly emerged males and workers were likely not inoculated at the same age. Indeed, workers developed in healthy colonies and were inoculated at the adult stage (most likely > 2 days old) for the establishment of infected microcolonies. However, males (most likely one day old) developed in infected microcolonies and ingested *Crithidia s*p. cells upon emergenceresulting in the infection of up to 90% of them (Gekière et al., 2021, *unpublished results*).. Second, while the difference in male fat body content between infected and uninfected microcolonies is unlikely to have arisen from a difference in brood care (i.e., no significant difference in pollen efficacy), we cannot rule out that infected workers displayed specific brood caring behaviour. For instance, they could have added peculiar nutrients or microorganisms to larval food from their hypopharyngeal and mandibular glands and/or stomach to prepare their offspring to face infection (e.g., addition of sterols, Svoboda et al., 1986). Such an increase in offspring fat body content through adapted larval feeding by workers could be interpreted as a trans-generational prophylactic behaviour. Indeed, enhanced fat body content has been assumed to correspond to a specific allocation of resources to counteract parasites by mounting an immune response (Brown et al., 2003). It would be interesting to test whether infected workers provide their larvae with specific central and specialised metabolites. We should however underline that the relationship between fat body content and immunity has become controversial since some results have been contradictory (e.g., Brown et al., 2000, Gekière et al., 2022a).

Although *Crithidia* sp. only showed mild effects in our experiment and in previous laboratory experiments (Brown et al., 2003; Goulson et al., 2018; Gekière et al., 2022a), it is important to keep in mind that interpretation of results observed in laboratory conditions must be interpreted with caution as controlled conditions are often not representative of natural constraints encountered in the field such as, for instance, predation, flight, and foraging. For example, infection by *Crithidia* sp. has been shown to impair pollen foraging (Shykoff & Schmid-Hempel, 1991; Otterstatter et al., 2005; Gegear et al., 2006), but such effects cannot be fully comprehended under laboratory conditions.

### Heather pollen quality: the case of flavonoids

Heather pollen harbours kaempferol flavonoids linked to one/two coumaroyl groups which are also linked to one/two hexosides (Gekière et al., *in prep*). Herein, we have shown that these heather flavonoids reduced the total offspring production, as indicated by a decreased pollen collection and a lower pollen efficacy, as well as a reduced mass of newly emerged males, thereby altering microcolony performance. Indeed, drone mass is known to impact flight distances, but also reproductive abilities, affecting the dissemination and reproductive success of bumble bee populations (Greenleaf et al., 2007; Amin et al., 2012). Such poor quality of heather pollen for the maintenance of buff-tailed bumble bee microcolonies has already been pinpointed (Vanderplanck et al., 2014). While it was partly attributed to its nutritional content (i.e., low concentration of amino acids and abundance of δ-7-avenasterol and δ-7-stigmasterol, Huang et al., 2011, Vanderplanck et al., 2014), our study demonstrated that specialised metabolites may also impact heather pollen quality, regardless of its nutritional content (i.e., central metabolites).

Both heather pollen and its flavonoids showed detrimental effects (i.e. reduction of offspring production, pollen efficacy). However, heather flavonoids seemed to induce a higher stress response than heather pollen as dilution behaviour was significantly higher in microcolonies fed the supplemented diet compared to those fed the control diet (i.e., mixing behaviour to mitigate unfavourable diet properties, Berenbaum & Johnson, 2015; Vanderplanck et al., 2018) while such a difference was not observed for microcolonies fed the heather diet. The reason of this discrepancy is not obvious, as both diets harbour the same flavonoids and should therefore lead to similar dilution behaviour. Two hypotheses could be proposed to explain this difference: (i) flavonoids were more bioavailable in the supplemented diets (outside pollen grains after the chemical extraction) and then more easily absorbed by the workers, which ultimately reduced the diet palatability (Wang et al., 2019); and (ii) as flavonoid extract was added to the control diet (i.e., willow pollen) that already contained flavonoids, the supplemented diet was richer in flavonoids than the other diets, reaching a threshold that ultimately reduced the diet palatability. Unfortunately, it is not possible to unravel these hypotheses without additional experiments. Another observation supporting the potential toxicity of heather flavonoids is the increase in fat body content in males emerging from microcolonies fed heather and supplemented diets compared to those emerging in microcolonies fed the control diet. Indeed, such an increase could be interpreted as a specific allocation of resources to the fat body for performing a detoxification activity (Li et al., 2019). In that way, flavonoid assimilation is known to induce the activation of defence mechanisms based on cytochrome P450 monooxygenase, a molecule that is highly active in the fat body (Scott et al., 1998). This increase in fat body content was not observed in workers, which could be explained by the different exposure to flavonoids during their life stages. Indeed, workers within microcolonies mainly fed on syrup, while males fed on pollen during their whole larval development and were then more exposed to specialised metabolites. Moreover, it is highly possible that sensitivity to pollen specialised metabolites is higher in larvae than in adults, as already demonstrated in honey bees (Lucchetti et al., 2018).

### The complex response of parasitised bumble bees to heather pollen and its flavonoids

Flavonoids were associated with an increase in parasite load, which has been also observed for other classes of specialised metabolites (Thorburn et al., 2015; Gekière et al., 2022a). Therefore, by contrast to our expectations based on previous studies (Baracchi et al., 2015, Koch et al., 2019), the detrimental effects of heather pollen flavonoids on bumble bees were not balanced by any therapeutic effect against the parasite *Crithidia* sp. These results suggest a potential additive effect between phytochemical and parasite stress as previously described (Thorburn et al., 2015), with the diet effect mostly overriding the parasite effect in bumble bees in the case of heather as already shown for the sunflower (Gekière et al., 2022a). The nutritional stress due to heather pollen feeding could then increase the effect of *Crithidia* sp. that could be more effective under stressful conditions (Brown et al., 2000; Brown et al., 2003).

We found that mortality in infected microcolony was lower in microcolonies fed the heather diet compared to those fed the control diet. Although infection was not shown to have a significant impact on mortality in microcolonies fed the control diet, heather pollen could then increase host tolerance to the parasite but this effect is unlikely due to its flavonoid content as mortality in infected workers fed the supplemented diet did not significantly differ from those fed the control diet. Regarding fat body content in newly emerged males, males that emerged in microcolonies fed the heather or supplemented diets displayed higher fat body content than those that emerged in microcolonies fed the control diet, but this diet effect was not significant anymore in infected microcolonies, probably because, as discussed before, the parasitic stress also increased fat body content in microcolonies fed the control diet.

## Conclusion

How heather pollen and its specialised metabolites impact the buff-tailed bumble bee, and how they modulate the interaction with its obligate gut parasite *Crithidia* sp. are complex questions given the diversity of specialised metabolites found in the floral resources of this species. Previous studies have found that heather nectar does not contain any flavonoids (Gekière et al., in prep) but protected the pollinator from its parasite *Crithidia* sp. through callunene activity (Koch et al., 2019). In this study, we found that the occurrence of flavonoids in heather pollen reduced its collection, as well as the bumble bee fitness. Moreover, heather pollen did not help to counteract the parasite but rather appeared to induce an additional stress that could potentially increase the parasite effect. Actually, our results complete the understanding of the bumble bee-heather-parasite relationship by underlining that heather pollen is not suitable to buff-tailed bumble bee performance and does not display any therapeutic effect. This study highlights the complexity of the plant-pollinator interaction by illustrating the distinct roles and effects of specialised metabolites found either in nectar or pollen. We strongly encourage the consideration of these two floral resources in future studies investigating the medicinal effects of plant species, especially when defining pollinator conservation strategies.

## Acknowledgements

We would like to thank D. Evrard and L. Marin for their help in colony and microcolony maintenance, as well as the numerous people that helped for microcolony dissection. We are very grateful to the Laboratory of Cellular Biology (UMons) and the Laboratory of Therapeutic Chemistry and Pharmacognosy (UMons) for their advice and the access to the device needed to the microscope analyses and the diet preparation. We also thank François Dittlo for providing us with heather pollen, as well as the family of the first author (I., C., C., and C. Tourbez) for their help.

## Authors’ contributions

Conceptualization, M.V.; chemical analyses and extraction, I.S. and P.G.; bioassays, C.T., with help from A.M. and A.G.; resources, D.M. and P.G.; writing—original draft preparation, C.T., with help from A.G. and M.V.; supervision, A.G. and M.V.; funding acquisition, D.M., P.G. and M.V. All authors have read and agreed to the published version of the manuscript.

## Data, script, and code information availability

Datasets and R script are avalaible on a Zenedo repository: https://doi.org/10.5281/zenodo.7804841

## Appendix A Supplemented diet

Total flavonoid content of heather bee pollen, as well as its associated dried extract, were analysed in triplicates by HPLC-MS/MS (triplicates of 20 – 40 mg) for quantification (expressed as quercetin equivalent, QE). Based on these analyses, the supplementation formula was established to have a similar amount of ethanol and willow pollen in candies for all diets, as well as flavonoid concentrations in the supplemented diet mimicking natural concentration found on average in bee pollen candies, namely 12.08 mg QE/heather candy on average (14.73 ± 1.69 mg QE/g for heather bee pollen) (Table S1). We found that heather bee pollen extract contained 40.63 ± 0.72 mg QE/g (209.79 g extract).

**Table S1.** Diet compositions.

| Diets | Ethanol<br>($\mu\text{L/g}$ candy) | Pollen<br>(g/g candy) | Flavonoids<br>(mg/g candy) |
| --- | --- | --- | --- |
| Control diet | 17.02 | 0.68 | 11.89 |
| Heather pollen-supplemented diet | 19.80 | 0.67 | 11.85 <sup>1</sup> |
| Heather diet | 20.50 | 0.82 | 12.08 |

<sup>1</sup> The concentration indicated for supplemented diets does not take into account the flavonoid concentration already present in the control diet (11.89 mg/g candy).

## Appendix B Variable estimates

**Table S2.**
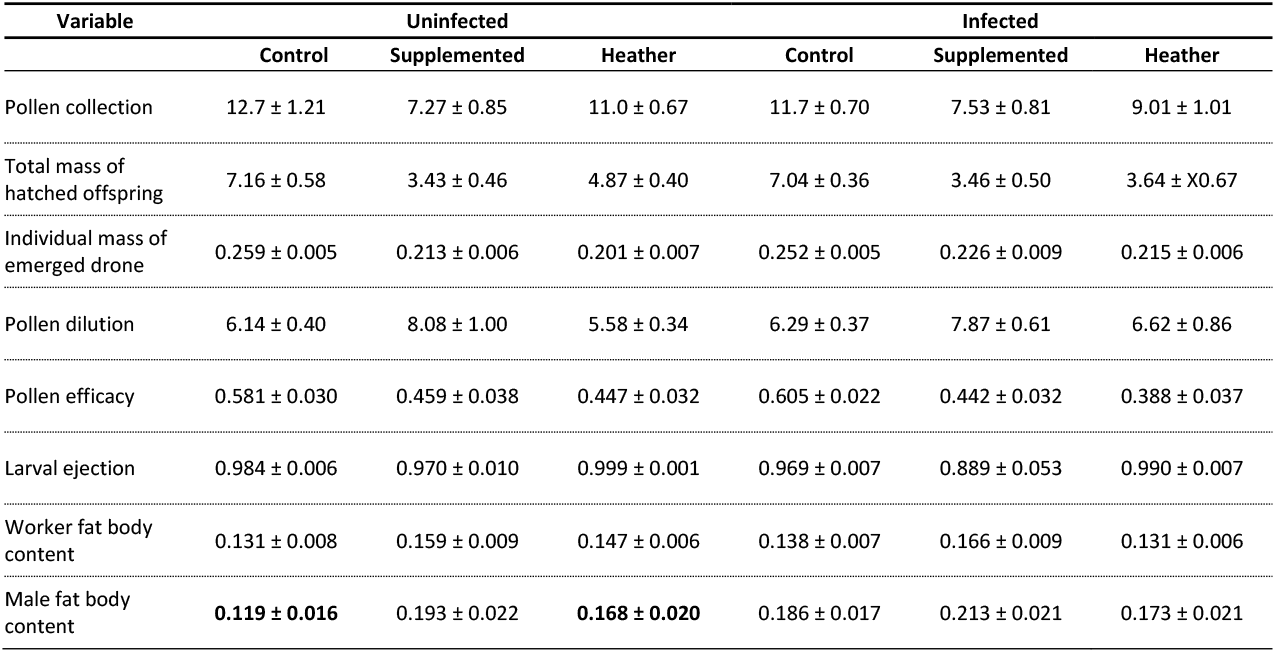
Mean ± standard error (SE) values of the variable used to describe parasite and diet effects.

